# Distinct streams for supervised and unsupervised learning in the visual cortex

**DOI:** 10.1101/2024.02.25.581990

**Authors:** Lin Zhong, Scott Baptista, Rachel Gattoni, Jon Arnold, Daniel Flickinger, Carsen Stringer, Marius Pachitariu

**Author notes:** these authors contributed equally (Carsen Stringer, Marius Pachitariu).

## Abstract

Representation learning in neural networks may be implemented with supervised or unsupervised algorithms, distinguished by the availability of feedback. In sensory cortex, perceptual learning drives neural plasticity, but it is not known if this is due to supervised or unsupervised learning. Here we recorded populations of up to 90,000 neurons simultaneously from the primary visual cortex (V1) and higher visual areas (HVA), while mice learned multiple tasks as well as during unrewarded exposure to the same stimuli. Similar to previous studies, we found that neural changes in task mice were correlated with their behavioral learning. However, the neural changes were mostly replicated in mice with unrewarded exposure, suggesting that the changes were in fact due to unsupervised learning. The neural plasticity was concentrated in the medial HVAs and obeyed visual, rather than spatial, learning rules. In task mice only, we found a ramping reward prediction signal in anterior HVAs, potentially involved in supervised learning. Our neural results predict that unsupervised learning may accelerate subsequent task learning, a prediction which we validated with behavioral experiments.

---

Many neurons in sensory cortical areas change their responses during learning and become more selective to task stimuli [1–13]. This has been interpreted as a possible basis for perceptual learning. A less common interpretation for neural plasticity is unsupervised learning, which has been proposed in multiple theoretical frameworks to describe how the brain learns from sensory experience, without the need for task labels or task feedback [14–19]. However, experimental evidence for such theories is limited to a few indirect observations. In primates, changes of neural tuning were observed in the inferotemporal cortex after repeated exposure to temporally-linked stimuli, even in the absence of rewards [20, 21]. In the mouse visual cortex, anticipatory responses to learned stimulus sequences have been interpreted as evidence for predictive coding, which is a form of unsupervised learning [22, 23]. Neural plasticity in the hippocampus has sometimes been interpreted as sensory compression [24, 25], which is also a type of unsupervised learning, though it has typically been linked to spatial rather than sensory representations [26, 27]. It is thus still not known how widely unsupervised learning may affect sensory neural representations.

Here we find that most of the neural plasticity in the visual cortex after task learning was replicated in mice with unsupervised exposure to the same visual stimuli. We found the only exception in anterior visual areas which encoded unique task signals potentially used for supervised learning. Below we describe a sequence of experiments designed to probe the roles of supervised and unsupervised learning across multiple visual computations, before and after learning.

## Both supervised and unsupervised training drive neural plasticity

Similar to previous work [10], we designed a visual discrimination task in head-fixed mice running through linear virtual reality corridors (Figure 1a). Mice had to discriminate between visual texture patterns in two corridors; these corridors were repeated in pseudo-random order. The visual patterns in each corridor were obtained as “frozen” crops from large photographs of naturalistic textures. For simplicity, we denote the stimuli as “leaf” and “circle”, even though other visual stimuli were also used in some mice. The visual stimuli were spatial frequency matched between categories to encourage the use of higher-order visual features in discrimination. A sound cue was presented at a random position inside each corridor and was followed by the availability of water in rewarded trials only (Figure 1a). After approximately two weeks of training (Figure 1b), mice demonstrated selective licking in the rewarded corridor in anticipation of reward delivery (Figure 1c,d, error bars on all figures represent s.e.m.). After learning, we introduced unrewarded test stimuli “leaf2” and “circle2”, which were different frozen crops of the same photographs. We then continued training with unrewarded “leaf2” until the mice stopped licking to this stimulus, at which point we introduced another test stimulus (“leaf3”) as well as spatially-shuffled versions of leaf1 (Figure 1b).

**Figure 1:**
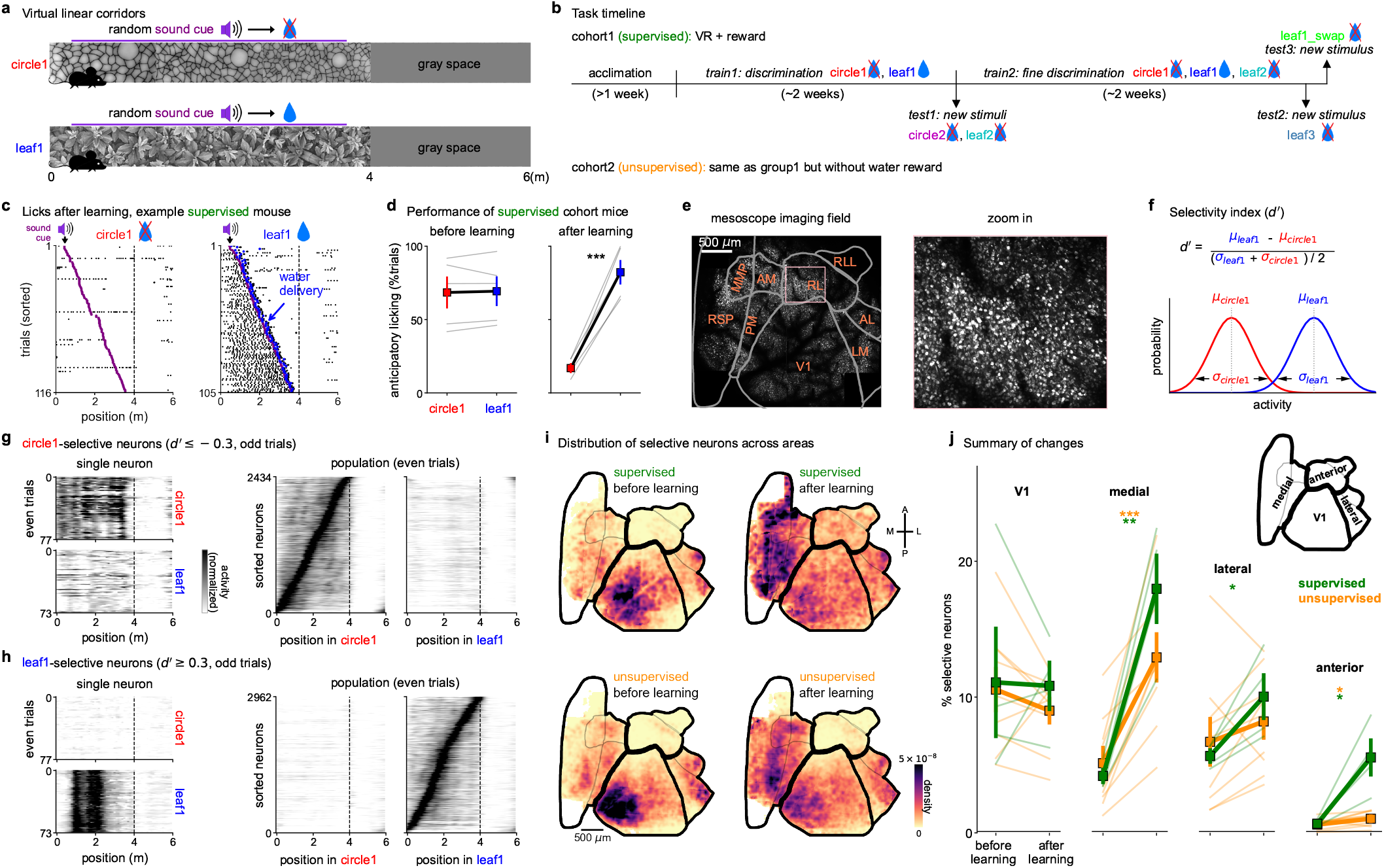
Plasticity in visual cortex after supervised and unsupervised training. **a**, Illustration of the virtual reality task with a sound cue at a random position in each corridor. Water is available after the sound cue in the rewarded corridor. **b**, Task training timeline in supervised mice. Mice in the unsupervised cohort experienced the same stimuli without water rewards. **c**, Lick distribution in an example mouse after task learning. Trials were sorted according to the sound cue position. **d**, Performance quantification of anticipatory licks before water is delivered (error bars on all figures represent s.e.m.; n=5 mice). **e**, Example field of view of the mesoscope and zoom in to illustrate cellular resolution. **f**, Selectivity index *d′* of neural responses inside the two corridors. **g**, Single-trial responses of a single circle1-selective neuron as well as the entire population from an example mouse. **h**, Same as **g** but for leaf1-selective neurons. **i**, 2D histogram of selective neuron distributions across the field of view, aligned to a map of visual areas. Top and bottom rows are supervised and unsupervised mice respectively. Left and right columns are before and after learning. **j**, Percentage of neurons with high-selectivity in each of four visual regions defined in the inset (n=4 supervised mice, n=9 unsupervised mice).

Mice in the unsupervised cohort also ran through the same corridors for similar periods of time, but did not receive water rewards and were not water-restricted.

Before and after learning, we recorded from large neural populations across many visual areas simultaneously using a two-photon mesoscope [28] (Figure 1e). We ran Suite2p on this data to obtain the activity traces from 20,547-89,577 neurons in each recording [29]. For each neuron, we computed a selectivity index d-prime (*d*′) using the response distributions across trials of each corridor, pooled across positions and for timepoints when the mice were running (Figure 1f). Neurons with relatively high d-prime (*d*′ ≥0.3 or *d*′≤-0.3) responded strongly at some positions inside the leaf1 or cicle1 corridor (Figure 1g,h). To see where these selective neurons were located, we generated 2D histograms of their position in tissue after aligning each session to a pre-calculated atlas of visual cortical areas (Figure 1i, Figure S1a,b, see also [30]). We found that, after learning, many selective neurons emerged in medial visual areas, encompassing regions PM, AM, and MMA, as well as the lateral part of the restrosplenial cortex [30] (Figure 1j). This region, which we will refer to as the “medial” visual region, showed similar changes in neural selectivity for both the supervised and unsupervised cohorts. These changes were also observed separately for neurons tuned to the leaf1 and circle1 stimuli respectively (Figure S1c). The lateral visual areas showed relatively little modulation by learning, and the anterior regions were only modulated in the supervised condition (see figure Figure 5 for more on this). We also observed some plasticity in V1, where the medial part showed a small but significant decrease in selectivity in the unsupervised cohort (Figure S1d), as well as some minor changes of selectivity when separated by stimulus (Figure S1c). However, the overall fraction of selective V1 neurons did not change much.

Thus, the distribution of neural plasticity across visual regions did not depend on task feedback or supervision.

## Visual, rather than spatial, representations after learning

It is possible that the neural plasticity we observed was due to spatial learning and navigation signals, which have been found to modulate firing rates even in the visual cortex [31]. Alternatively, the neural plasticity might be due to adaptation to the visual statistics of the natural images we presented [14]. To distinguish between a spatial and a visual plasticity hypothesis, we next introduced two unrewarded test stimuli, leaf2 and circle2, which contained similar visual features to leaf1 and circle1 respectively, but were arranged in different spatial configurations (Figure 2a). We found that mice only licked to leaf2 and not circle2, likely due to their visual similarities with the trained stimuli (Figure 2b,c).

**Figure 2:**
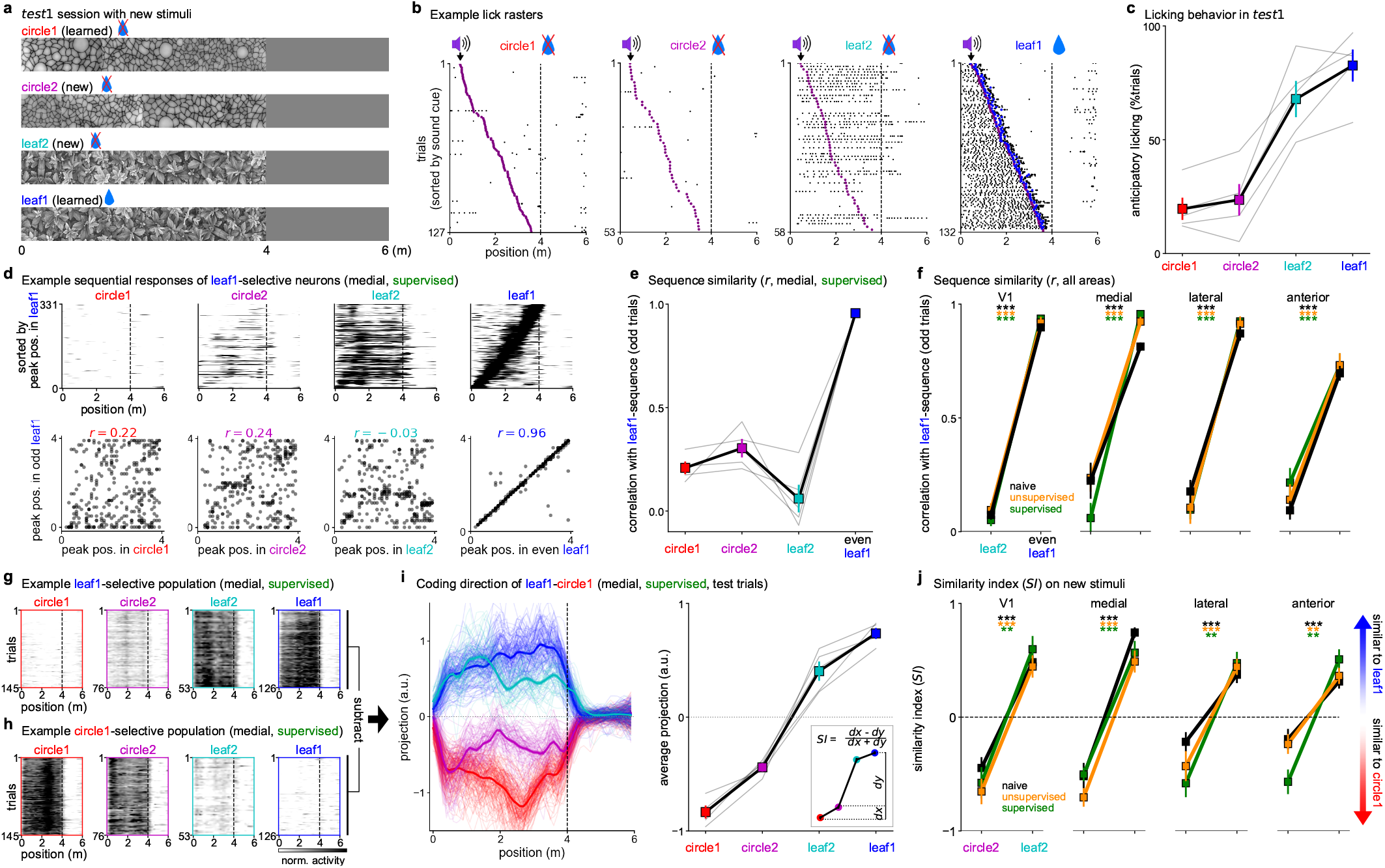
Comparing visual and spatial coding on test stimuli. **a**, Stimuli in the *test1* session (see timeline in Figure 1b). **b**, Lick rasters for an example mouse (blue dots represent reward delivery). **c**, Anticipatory licking behavior in *test1* (n=5 mice). **d**, (top) Example neural responses from the medial region in a mouse from the supervised cohort, sorted by preferred position in the leaf1 corridor on held out trials. (bottom) Scatter plots of preferred positions in the leaf1 corridor versus all other corridors. **e**, Correlation from **d** (bottom) for each mouse in the supervised cohort (n=5 mice). **f**, Same as **e** but only for the leaf2 and leaf1 corridors, shown across regions and for naive (n=5 mice, 6 sessions), supervised (n=5 mice) and unsupervised (n=7 mice). **g**,**h**, Example leaf1- and circle1-selective neurons respectively, shown as a population average over trials, **i**, Projections onto the coding direction, defined as the difference between the leaf1- and circle1-selective populations in **gh**; (left) timecourse, (right) average over trial and (inset) definition of similarity index for circle2. **j**, Similarity index for new stimuli.

The spatial plasticity hypothesis suggests that neurons would fire in a similar sequence to learned and new exemplars of the same category. We tested this directly by sorting neurons according to their sequence of firing in the leaf1 corridor. This sorting did not induce similar sequences for the leaf2 corridor in the medial regions (Figure 2d). The neural sequences in leaf1 and leaf2 were in fact uncorrelated in all regions, suggesting a non-spatial coding scheme (Figure 2e,f). Furthermore, the sequence correlations did not match the behavioral strategy of the mice (Figure 2c,e). Similar results were found when analyzing the sequences of the circle1-selective neurons (Figure S2a,b).

The visual plasticity hypothesis suggests that the statistics of visual features (i.e. “leafiness”) are learned regardless of where the features occur in the corridor. To test this, we designed an analysis which uses the top selective neurons of each familiar corridor to create a coding direction axis [32] (Figure 2g,h). Projections of neural data on the coding direction were well-separated on test trials of the familiar stimuli, and also on trials of the new leaf2 and circle2 stimuli (Figure 2i, Figure S2c-e). To quantify this separation, we defined a similarity index (*SI*) computed from the projections on the coding axis. The similarity index clearly distinguished between corridors of different visual categories, in all visual regions, in both supervised and unsupervised mice as well as in naive mice (Figure 2j). Thus, the coding direction readout of neural activity matched the behavioral strategy of the mice (Figure 2c,i), suggesting that the brain areas we consider use visual rather than spatial coding.

## Novelty responses and neural orthogonalization

While the neurons selective to familiar stimuli also responded to the new stimuli, they were by far not the most responsive neurons to the new stimuli. Selecting neurons by d-prime between leaf2 and circle1, we found a large population of leaf2-selective neurons in V1 and the lateral visual areas (Figure 3a, see also Figure S3a for circle2-selective neurons). The responses of leaf2-selective neurons adapted substantially after an additional week of training with the leaf2 stimulus (Figure 3a,b). In contrast, leaf1-selective neurons showed little or no change after this additional learning phase (Figure S3b). Thus, V1 and lateral HVAs appear to be responding to the stimulus novelty, as reported in some previous studies [33], and the responses were similar in the supervised and unsupervised cohorts.

**Figure 3:**
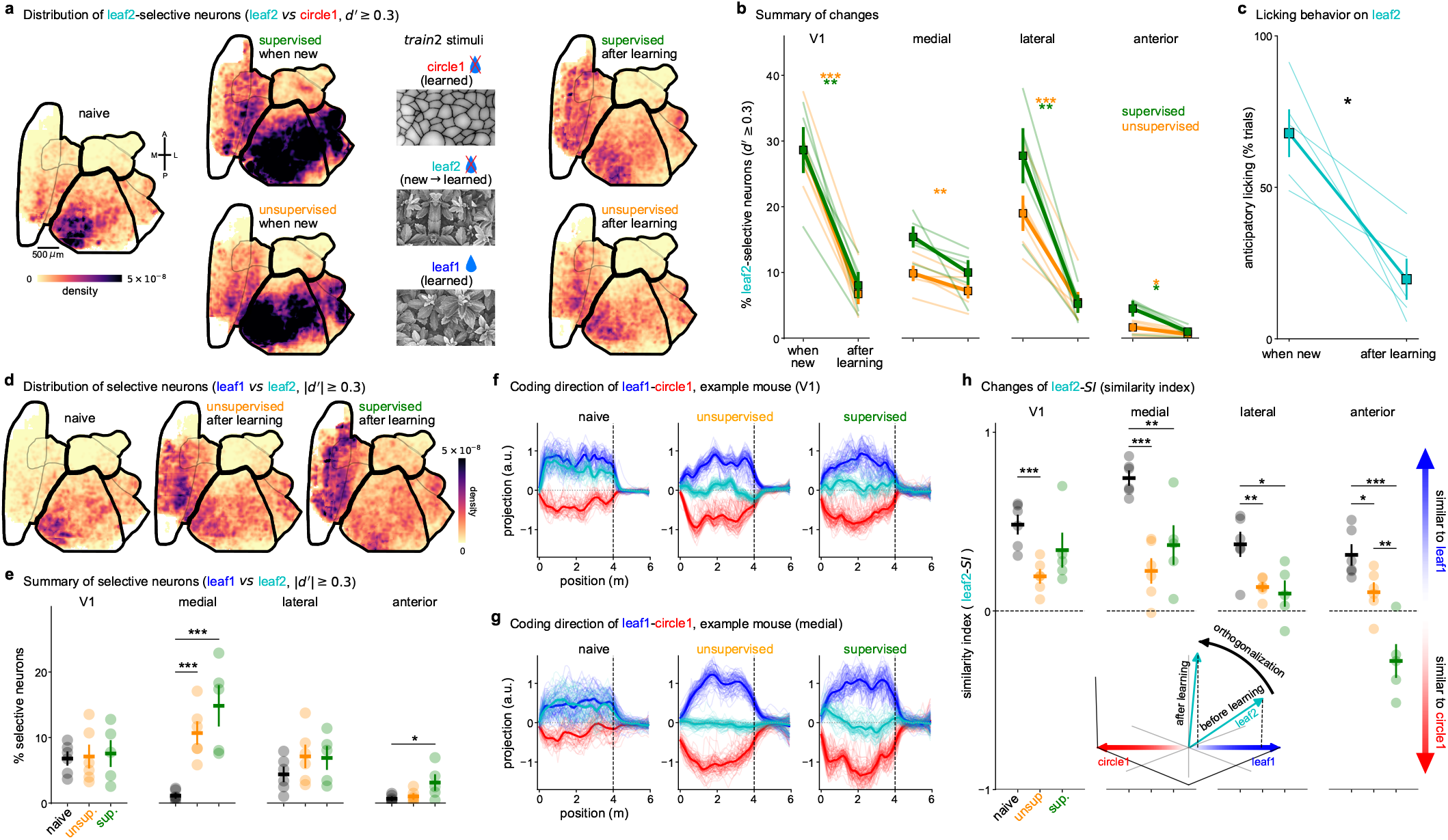
Responses to novel and adapted stimuli and neural orthogonalization. **a**, Distribution of neurons with high *d′* between the leaf2 and circle1 corridors in supervised and unsupervised mice, either when new or after learning, as well as in fully naive mice. **b**, Summary of distribution changes in **c** across regions (n=5 supervised, n=6 unsupervised mice). **c**, Licking behavior to leaf2 when new and after learning (n=5 mice). **d**, Distribution of neurons with high *d′* between leaf1 and leaf2 in supervised and unsupervised mice after training with leaf2, as well as in fully naive mice. **e**, Summary of **d** across regions (n=5 naive mice (6 sessions), n=5 supervised mice, n=6 unsupervised mice). **f**,**g**, Example projections on the coding direction between leaf1 and circle1 in V1 (**f**) and the medial region (**g**). **h**, Similarity index (from Figure 2i) for neural responses to leaf2 in supervised (n=5) and unsupervised (n=6) mice after training with leaf2, as well as in fully naive mice (n=5 mice, 6 sessions). Bottom inset shows schematic of orthogonalization effect observed. Neural vectors are referenced with respect to the center of the leaf1-circle1 axis.

The task mice eventually stopped licking to leaf2 (Figure 3c), thus indicating strong discrimination between leaf1 (rewarded) and leaf2 (unrewarded). We hypothesized that this behavior may be accompanied by changes in neural discrimination, similar to the changes after mice learned the distinction between leaf1 and circle1 (Figure 1j). To test this, we selected neurons based on their *d*′ between leaf1 and leaf2, and compared the fraction of tuned neurons across areas in naive mice as well as in the supervised and unsupervised cohorts. Again we observed a substantial increase in selectivity in the medial HVAs, in both the supervised and the unsupervised cohorts (Figure 3d,e).

Because leaf1 and leaf2 neural representations were similar in naive mice (Figure 2j), we hypothesized that the fine behavioral discrimination between leaf1 and leaf2 stimuli requires orthogonalization of their neural representations [27, 34]. We tested this by comparing the projections of leaf2 onto the leaf1-circle1 coding direction (Figure 3f,g). Compared to naive mice, this projection was reduced after both supervised and unsupervised training, across all visual regions but most strongly in the medial HVAs (Figure 3h). Thus, the responses to leaf2 became orthogonal to the leaf1-circle1 axis, even in the unsupervised mice in which leaf1 and leaf2 had the same valence. Altogether these observations describe the complex dynamics of the representation of a new stimulus (leaf2) as it becomes familiar, in both supervised and unsupervised conditions.

## Visual recognition memory

By this stage in the training, mice had been exposed to the leaf1 stimulus for ∼ 4 weeks, and the supervised cohort had learned to distinguish it from leaf2. We hypothesized that a more detailed representation of leaf1 emerged to support the visual recognition memory of this stimulus. To test this, we first introduced a new exemplar of the leaf category (“leaf3”, Figure 4a). Mice withheld licking to this stimulus, similar to their behavior on the unrewarded, trained leaf2 stimulus (Figure 4b). This behavioral choice mirrored a change in neural tuning properties: the neural projections of leaf3 trials onto the coding axis of leaf1-leaf2 were strongly biased towards the leaf2 direction (Figure 4c). This asymmetry was present in both the supervised and unsupervised cohorts, and it was not present in naive mice (Figure 4d,e). All visual regions behaved in this manner, except the anterior region where the coding asymmetry was more pronounced for supervised compared to unsupervised training, and the representation of the “control” stimulus (circle1) was also biased towards the leaf2 axis. As we will see in the next section, this may be due to the reward valence coding in anterior HVAs. To further test the visual recognition memory for the leaf1 corridor, we introduced two new corridors with spatially-swapped portions of leaf1 (Figure 4f and Figure S4a). We reasoned that these new corridors should disrupt the mice if they had memorized only the beginning of the corridor, or if they had used a purely spatial-based memorization strategy. We found no such disruption, with the mice licking in the swapped corridors at levels comparable to the leaf1 corridor (Figure 4g). Thus, the swapped corridors were still recognized as visually similar to the rewarded leaf1 corridor, as opposed to being recognized as a different exemplar of the leaf category (like leaf2 or leaf3). This behavior was supported by a neural coding strategy that tied neural response vectors to their respective visual stimulus locations (Figure 4h, Figure S4b-d). This visual encoding was present in all areas in both the supervised and unsupervised training conditions, and it was also present before training in V1, as expected for a purely visual representation (Figure 4i). The medial region did not appear to strongly encode a sequence before learning (Figure 4h,i), but this result may have been due to lower SNR from an overall weaker stimulus tuning before learning (Figure 1j).

**Figure 4:**
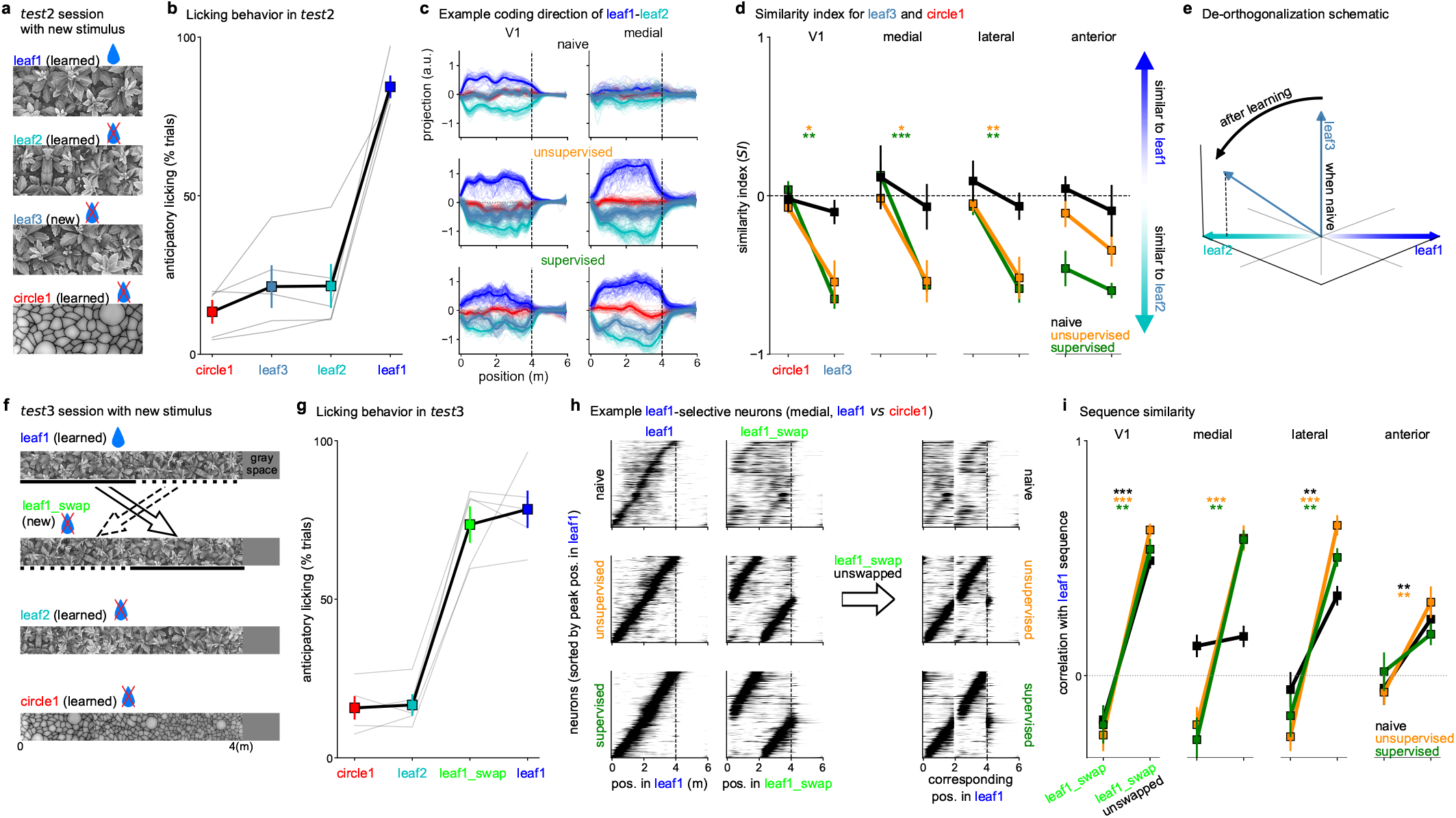
Visual recognition memory becomes exemplar-specific after extended training. **a**, Stimuli in the *test2* session (see timeline in Figure 1b). **b**, Anticipatory licking behavior in *test2* (n=5 mice). **c**, Example projections of neural data onto the coding direction of leaf1 vs leaf2. **d**, Similarity index (from Figure 2i) of leaf3 and circle1 stimuli for the leaf1 vs leaf2 coding direction (n=3 naive mice, 5 sessions; supervised n=5 mice; unsupervised n=6 mice). **e**, Schematic of observed “de-orthogonalization” effect, where an initially symmetric projection of leaf3 becomes asymmetric and more similar to the leaf2 neural vector. Neural vectors are referenced with respect to the center of the leaf1-leaf2 axis. **f**, Stimuli in the *test3* session (see timeline in Figure 1b). **g**, Anticipatory licking behavior in *test3* (n=3 mice, 5 sessions). **h**, Example leaf1-selective neurons (medial region) during leaf1 and swap trials, sorted by responses in the leaf1 corridor for naive, supervised and unsupervised mice. Also shown (right) are the responses on swap trials after reversing the swap manually (“unswapping”). **i**, Average correlation in position preference between leaf1 and the swap as well as the swap (unswapped) responses, shown for all three groups of mice across regions (n=3 naive mice, 10 sessions; n=3 supervised mice, 5 sessions; n=4 unsupervised mice, 8 sessions).

Thus, for both the leaf3 and leaf1-swap stimuli, behavioral responses were linked to the patterns of neural responses during supervised training, but these patterns were mostly present after unsupervised training as well.

## A reward prediction signal in anterior HVAs

Having found multiple similarities between the supervised and unsupervised conditions, we next asked whether a more targeted analysis could reveal differences. Since we did not know in advance what to look for, we used Rastermap, a visualization method for large-scale neural responses [35]. Rastermap reorders neurons across the y-axis of a raster plot, so that nearby neurons have similar activity patterns. Inspected in relation to task events, Rastermap can reveal single-trial sequences of neural activity tied to corridor progression, as well as other signals that may be related to task events like rewards and sound cues (Figure 5a). One of the signals we found with Rastermap corresponded to a neuronal cluster that turned on specifically in the leaf1 corridor but not in the circle1 corridor, and was turned off by the delivery of reward (Figure 5a,b). These neurons were distributed in the anterior HVAs (Figure 5c), and their activity was strongly suppressed by the delivery of reward (Figure 5d), similar to a reward prediction signal [36, 37].

**Figure 5:**
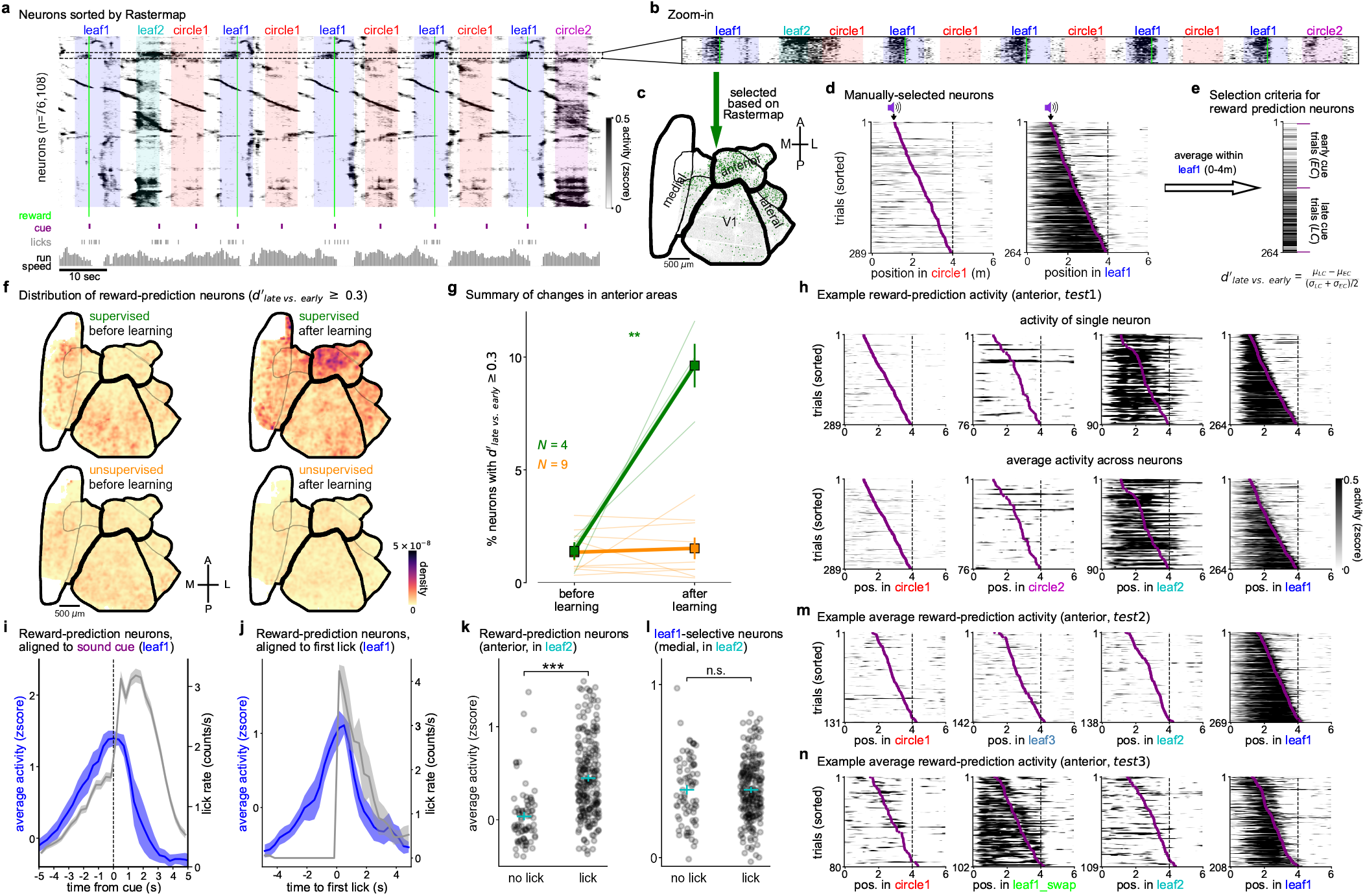
A reward prediction signal in supervised training only. **a**, Raster plot of *>*50,000 simultaneously-recorded neurons, over a period of approximately 2.5 minutes, with behavioral annotations. Rastermap was used to sort neurons across the y-axis. **b**, Zoom-in of a vertical segment in Rastermap corresponding to a group of neurons active in the leaf1 corridor. **c**, Spatial distribution of selected neurons. **d**, Population average response for the neurons selected in **b** across trials of circle1 and leaf1, sorted by sound cue position. **e**, A discrimination index 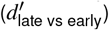 was constructed to compare trial-averaged neuron responses between trials with early vs late rewards. **f**, Distribution of reward prediction neurons 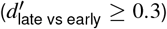 in supervised and unsupervised mice, before and after learning. **g**, Proportion of reward prediction neurons in the anterior region. **h**, Responses of a single selective neuron (top) or the entire population (bottom) across trials in the *test1* session. **i**, Reward prediction neuron activity and lick rates in the leaf1 corridor, aligned to the sound cue in the *test1* session, averaged across mice (n=5). **j**, Same as **i** but aligned to the first lick in the corridor, for trials where the first lick was after 0.5m (n=4 mice). **k**, Average activity of reward prediction neurons inside leaf2 corridor on trials with or without licks in the *test1* session, pooled across all mice (n=5). **l**, Same as **k** but for the leaf1-selective neurons in the medial regions (defined like in Figure 1h). **m-n**, Same as **h** (bottom) for the *test2* and *test3* sessions.

Having found a putative task-related population with Rastermap, we next quantified the task modulation at the single neuron level across mice. For this, we developed an index 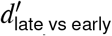 which compares neural activity on trials where the reward is delivered early to trials where it is delivered late (Figure 5e). Selective neurons 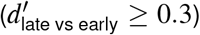 were distributed primarily in anterior areas in supervised mice after training (Figure 5f,g, Figure S5a). We did not observe a similar population when selecting neurons based on circle1 trials with a similar process (Figure S5b,c). Some single neurons in the anterior area with high 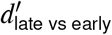 encoded the reward prediction signal robustly at the single trial level, and their response generalized to the new exemplar leaf2 of the rewarded visual category, similar to the population average of selective neurons (Figure 5h).

One possibility is that the reward prediction signal directly reflects licking behavior, because it was present only in the supervised cohort. We do not think this was the case, because 1) the reward prediction signal was strongly suppressed after the cue, whereas licking increased dramatically (Figure 5i and Figure S5d-f); 2) the reward prediction signal started ramping up several seconds before the first lick (Figure 5j). These dynamics are indicative of a reward expectation, especially because the neural prediction signal was higher on leaf2 (unrewarded) trials where the mouse licked compared to trials where it did not (Figure 5k). This distinction was only found in anterior HVAs, and not for example in the medial population we described above (Figure 5l).

The reward prediction signal continued to follow the dynamics of the behavior itself over the course of training. After training with leaf2, the reward prediction signal was suppressed during the leaf2 corridor, and was also absent in the new leaf3 corridor (Figure 5m). Finally, the reward prediction signal was present in the swapped leaf1 corridor, again indicating that this signal correlates with the expectation of reward (Figure 5n).

## Unsupervised pretraining improves perceptual learning in mice

Next we tested the potential function of the neural plasticity after unsupervised training. We hypothesized that this plasticity might allow animals to learn a subsequent task faster, similar to how unsupervised pretraining helps artificial neural networks to learn supervised tasks faster, and similar to previous maze learning experiments [38]. We thus ran a behavioral study in which one cohort of mice (“no pretraining”) was trained similarly to the task mice above, while a second cohort (“unsupervised pretraining”) first underwent 10 days of VR running without rewards (Figure 6a, Figure S6a). Compared to our original task, we simplified reward learning by restricting reward delivery to the second half of the reward corridor (the “reward zone”) and removing the sound cue (Figure 6b).

**Figure 6:**
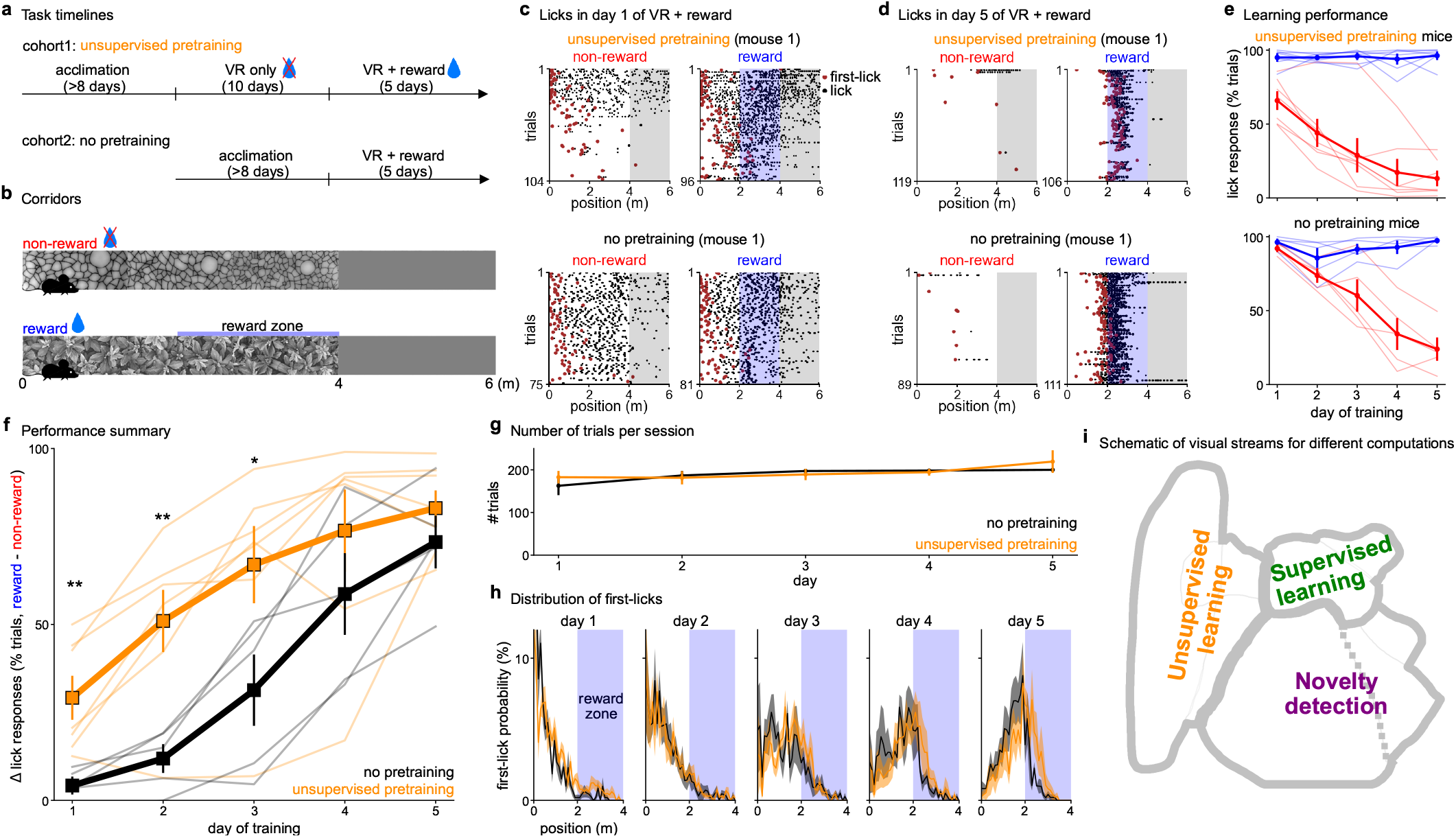
Unsupervised pretraining accelerates subsequent task learning. **a**, We trained two new cohorts of mice with or without a pretraining step of running through the VR without rewards for 10 days. Both cohorts were trained in the task for five days. **b**, Task structure (sound cue was removed and rewards were delivered deterministically in the second half of the reward corridor). Mice had to lick to obtain reward, except on the first day when rewards were delivered passively at the end of the corridor. Each mouse was trained on a separate combination of wall textures from among four stimuli (Figure S6a). **c**, Licking of example mice from each cohort on the first day of training. **d**, Same as **c** for the last day of training. **e**, Average lick responses across days for both cohorts of mice (n=7 cohort1 mice, n=5 cohort2 mice). **f**, Performance summary (difference in lick responses) across days for both cohorts. **g**, Numbers of trials per day for both cohorts. **h**, Distribution of first licks across days for both cohorts. **i**, Schematic of putative visual streams for different learning algorithms and visual computations.

Mice with unsupervised pretraining generally learned the task much faster. For example, one mouse started the first day of task training by licking indiscriminately in both corridors, but stopped licking in the non-reward corridor after ∼ 50 trials of each corridor (Figure 6c). By the fifth day of training, this mouse was selectively licking only at the beginning of the reward zone (Figure 6d). In contrast, mice without pretraining did not learn to distinguish the two corridors on the first day of training (Figure 6c,d). After five days, both cohorts reached a high discrimination performance, but the unsupervised pretrained cohort learned faster (Figure 6e,f). Further, most of the learning improvements happened within session (Figure S6b). The improved discrimination ability of the pretrained mice was not due to differences in behavior during task learning: both cohorts ran similar numbers of trials, and licked at similar positions in the two corridors (Figure 6g,h).

## Discussion

Here we showed that unsupervised learning has a substantial effect on neural representations in cortical visual areas, and helps mice learn a supervised task faster. The main region for unsupervised learning may be the medial HVAs, as these areas contained emergent representations that strongly discriminated the learned stimuli and emerged with or without task training (Figure 6i). However, the medial population did not represent trial-to-trial variability in mouse decision-making, while anterior HVAs did (Figure 5k,l). The reward prediction signal in anterior HVAs may be required for supervised or reinforcement learning. Finally, stimulus novelty was represented in the responses of neurons in V1 and lateral HVAs (Figure 6i).

Our results can be related to other reports of neural plasticity in sensory cortices. In the visual cortex, learning can result in more neurons that discriminate the learned stimuli [1, 10, 13], neurons that discriminate the stimuli better [2, 9, 12], or neurons that respond more to the learned stimuli [3, 6, 8]; learning can also add a context-dependence to the visual tuning of neurons [31], and it can orthogonalize stimulus representations in V1 [34]. Such changes have typically been interpreted as consequences of task learning because they correlate well with task performance. However, our results suggest that these changes would have happened even without task training.

Our results can also be compared to those in the hippocampus. While we have shown that the cortical representations are visual, rather than spatial in nature, it is possible that hippocampal representations also inherit some visual properties from their inputs [39– 41]. Further distinguishing between spatial and visual learning in the same circuits could be a promising direction of future research. Another promising direction could be to relate the unsupervised plasticity we observed here to the many classical theories and models of unsupervised learning [14–18] as well as to modern approaches like self-supervised learning [19, 42–45].

## Acknowledgments

This research was funded by the Howard Hughes Medical Institute at the Janelia Research Campus. From the Vivarium, we thank Jim Cox, Crystall Lopez, Anne Kuzspit, Miriam Rose, Michalis Michaelos, Gillian Harris, Sarah Lindo, and their respective teams for animal breeding, husbandry, surgeries, and behavioral training support. From JeT, we thank Alex Sohn, Tobias Goulet, Dmitri Tsyboulski and Steven Sawtelle for help with rig maintenance and upgrades. From MBF Bioscience we thank Georg Jaindl, Mitchell Sandoe and Boris Djiguemde for scanimage support. We thank Sandro Romani, Vivek Jayaraman, and Nelson Spruston for helpful discussions about the work.

## Author contributions

LZ and MP designed the study. LZ performed the imaging experiments. LZ, SB, and RG performed behavioral experiments. LZ and JA designed and maintained the behavioral apparatus. DF maintained and improved the mesoscope. LZ, CS, and MP performed data analysis. LZ, CS, and MP wrote the manuscript with input from all authors.

## Data availability

The data will be made available upon publication in a journal.

## Code availability

Analysis code will be made available upon publication in a journal.

## Methods

All experimental procedures were conducted according to IACUC, and received ethical approval from the IACUC board at HHMI Janelia Research Campus.

## Experimental methods

### Animals

We performed 79 recordings in 15 mice bred to express GCaMP6s in excitatory neurons: TetO-GCaMP6s x Emx1-IRES-Cre mice (available as RRID:IMSR JAX:024742 and RRID:IMSR JAX:005628) [46]. These mice were male and female, and ranged from 2 to 11 months of age. Mice were housed in reverse light cycle, and were pair-housed with their siblings before and after surgery. The mice had a running wheel in their cage, as well as corncob bedding with Nestlets. During training and imaging periods, we replaced the running wheel with a tube, to potentially motivate the mice to run longer while headfixed. Due to the stability of the cranial window surgery, we often use the same mice for multiple experiments in the lab: two of the mice were used in reference [47].

We also used 12 C57 mice for behavior-only experiments. These mice were only implanted with a headbar, not a cranial window.

### Surgical procedures

Surgeries were performed in adult mice (P35–P333) following procedures outlined in reference [48]. In brief, mice were anesthetized with Isoflurane while a craniotomy was performed. Marcaine (no more than 8 mg/kg) was injected subcutaneously beneath the incision area, and warmed fluids + 5% dextrose and Buprenorphine 0.1 mg/kg (systemic analgesic) were administered subcutaneously along with Dexamethasone 2 mg/kg via intramuscular route. For mice with cranial windows, measurements were taken to determine bregma-lambda distance and location of a 4 mm circular window over V1 Cortex, as far lateral and caudal as possible without compromising the stability of the implant. A 4+5 mm double window was placed into the craniotomy so that the 4mm window replaced the previously removed bone piece and the 5mm window lay over the edge of the bone. After surgery, Ketoprofen 5mg/kg was administered subcutaneously and the mice were allowed to recover on heat. The mice were monitored for pain or distress, and Ketoprofen 5mg/kg was administered for 2 days following surgery.

### Imaging acquisition

We used a custom-built 2-photon mesoscope [28] to record neural activity, and ScanImage [49] for data acquisition. We used a custom online Z-correction module (now in ScanImage), to correct for Z and XY drift online during the recording. As described in reference [48], we used an upgrade of the mesoscope that allowed us to approximately double the number of recorded neurons using temporal multiplexing [50].

The mice were free to run on an air-floating ball. Mice were acclimatized to running on the ball for several sessions before training and imaging.

### Visual stimuli

We showed virtual reality corridors to the mice on three perpendicular LED tablet screens which surrounded each mouse (covering 270 degrees of their visual field of view). To present the stimuli, we used PsychToolbox-3 in MATLAB [51]. The virtual reality corridors were each 4 meters long, with 2 meters of gray space between corridors. The corridors were shown in a random order. The mouse moved forward in the virtual reality corridors by running. Running was detected using an optical tracking sensor placed close to the ball.

The virtual reality corridors were created by concatenating 4 random crops from one of four large texture images: circle, leaf, rock, and brick.

### Water restriction procedure

Water restriction procedures were conducted according to IACUC. During the VR + reward training, animals received an average of 1 mL water per day (range 0.8-1.2 mL depending on health status and behavioral performance). Before reaching 1 mL water per day after the initiation of the restriction procedure, we gradually reduced the water amount from 2 mL per day, to 1.5 mL per day until finally to 1 mL per day. The behavior-only mice were water restricted for 5 days right before the VR + reward training. Once the animals finished the VR + reward training session, the remaining water (0.8-1.2 mL minus the amount received during experiment) was provided 0.5 hr after the training. During the whole water restriction period, the body weight, appearance, and behaviors were monitored using a standard quantitative health assessment system [52].

### Water reward delivery and lick detection

A capacitance detector was connected with the metal lick port to detect licking. Mice received a drop of water (2.5 uL) if they correctly licked inside the reward corridor. In day 1 of VR + reward training, we always delivered the water passively (passive-mode) so that the mice could get used to acquiring reward when stimuli were present. For all the behavior-only mice (Figure 6) and some of the imaging mice (Figures 1-5), we switched to active-reward mode after day 1 so that the mice had to lick within the reward zone in order to trigger the water delivery. For some of the imaging mice (Figures 1-5), we kept using the passive-mode but added a delay (1s or 1.5s) between the sound cue and reward delivery. Given that mice started licking as soon as they entered the corridor and until they received the water, adding a delay *vs*. active-reward mode did not change how the mice behaved (Figures 1-5).

### Behavioral training

All animals were handled via refined handling techniques for at least 3 days prior to being acclimated to head-fixation on the ball. Animals were acclimated gradually (0.5 to 1 hr per day) on the ball over at least 3 days until they could be head-fixed without exhibiting any signs of distress. Then, animals began a running training regiment (1 hr per day) which lasted for at least 5 days to ensure they could run smoothly and continuously on the ball before being exposed to the closed-loop virtual linear corridor. For water restricted mice, we trained them for two days to get used to acquiring water from the spout when no stimulus was presented, before the VR + reward training. For the unsupervised pretraining group of mice (Figure 6), learning to get reward from the spout was carried out on the last two days of unsupervised pretraining, after the VR session to avoid the associative learning between stimuli and rewards. For the group without pretraining (Figure 6), learning to get reward from the spout was similarly carried out after the running training session on the last two days of running training.

For the behavior-only experiment, all animals started training in VR + reward training on a Monday and continued training for exactly 5 days. This ensured a consistent training schedule during the critical learning period.

For imaging mice, we consider a lick response if the mouse licks at least once inside the corridor but before the reward delivery. For the behavior-only mice (Figure 6) we consider a lick response if the mouse licks inside the reward zone (second half of the corridor) at least once.

## Data analysis

For analysis we used Python 3 [53], primarily based on numpy and scikit-learn [54, 55], as well as rastermap [35]. The figures were made using matplotlib and jupyter-notebook [56, 57].

### Processing of calcium imaging data

Calcium imaging data was processed using Suite2p [29], available at www.github.com/MouseLand/suite2p. Suite2p performs motion correction, ROI detection, cell classification, neuropil correction, and spike deconvolution as described elsewhere [58]. For non-negative deconvolution, we used a timescale of decay of 0.75 seconds [59].

### Neural selectivity (d-prime)

To compute the selectivity index d-prime (*d*′), illustrated in Figure 1f, we only selected data points inside the 0-4m region of the corridors where the textures were shown. We excluded the data points in which the animal was not running, so that all data points included for calculating the selectivity index come from similar engagement/arousal levels of the mice. We first calculated the means (*μ*_1_, *μ*_2_) and standard deviations (σ_1_, σ_2_) of activities for any two corridors, then computed the *d’*. The criteria for selective neurons was |*d*′| ≥ 0.3:

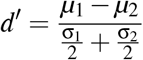

To make the density plots across the cortex (e.g. Figure 1i), we computed 2D histograms for each session based on the selective neurons in that session. We then applied a 2D Gaussian filter to this matrix and divided by the number of total recorded neurons in that session to get a density map for each mouse. Before averaging the density maps across mice, we assigned NaN to areas where no neurons were recorded. This ensured no underestimation on the density within areas where not all mice have neurons recorded.

For sequence similarity analyses (Figure 2, Figure 4, Figure S2, Figure S4), we used half of leaf1 and circle1 trials (train trials) to compute the selectivity index *d*′and we selected neurons based on the criteria |*d*′|≥0.3. We then split the other half of leaf1 and circle1 trials (test trials) into odd *vs*. even trials to compute spatial tuning curves for odd and even trials separately for each selective neuron. From these spatial tuning curves, we used the position with the maximal response as the preferred position for each neuron. To compute tuning curves for other stimuli such as leaf2, circle2 and swap (which were not used to find selective neurons), we split all trials into odd and even trials. The preferred positions of the same neurons in different corridors or in odd vs. even trials were used to compute a correlation coefficient (*r*).

*Coding direction and similarity index*

To compute the coding direction (Figure 2, Figure 3, Figure 4, Figure S2, Figure S4), for example in leaf1 vs circle1, we first chose leaf1- and circle1-selective neurons based on their *d*′ from the train trials (using the top 5% selective neurons each to leaf1- and circle1-, same as the sequence similarity analysis). Then we normalized the neural activity **r** for each neuron by subtracting the baseline response in the gray portion of the corridor *μ*_*gray*_, and dividing by the average standard deviation of the neuron’s responses in each corridor:

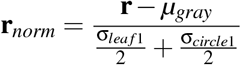

We then computed the mean normalized activity ***μ***_*leaf* 1_ of leaf1-selective neurons and the mean normalized activity ***μ***_*circle*1_ of circle1-selective neurons at each position in each corridor. The coding direction 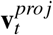 on a given trial *t* was defined as the difference

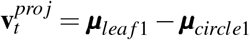

Note this is equivalent to assigning weights of 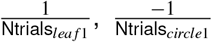 and 0 respectively for positively-selective, negatively-selective and non-selective neurons, and using those weights as a projection vector for the neural data. We investigated the coding direction always on test trials not used for selecting neurons, either from held-out trials of leaf1 and circle1, or for trials with other stimuli. We averaged the responses across each trial type: 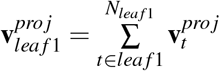 (e.g. Figure 2i, left).

Average projections for each trial type were computed by averaging these projections within the texture area (0-4m) (e.g. Figure 2i, right), denoted as 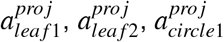 and 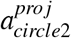. We then defined the similarity index *SI* on a per-stimulus basis, for example for leaf2, as:

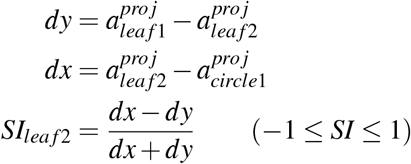

which is quantified in Figure 2j. We also computed the coding direction for different sets of selective neurons, e.g. leaf1 vs leaf2, and then computed the similarity indices for leaf3 and circle1 (Figure 4c,d).

### Reward-prediction neurons

Reward-prediction neurons were either selected using the clustering algorithm Rastermap (Figure 5a-d) or using a *d*′ criterion (Figure 5e-n). Using Rastermap, we selected the reward-prediction neurons based on their special firing patterns of only responding inside the rewarded corridor and specifically before reward delivery. Using *d*′, we first interpolated the the neural activity of single neurons based on their position inside the corridor and constructed a matrix (trials by positions). Only the leaf1 trials (rewarded for supervised mouse cohort, and unrewarded for the unsupervised cohort) were chosen and divided into early-cue trials *vs*. late-cue trials based on the sound cue position inside the corridor. We used cue position instead of reward position because the sound cues were played in each corridor at a random position, with or without reward, and these sound cue positions were highly correlated with reward positions in the rewarded corridor (Figure 1c). We then calculated the 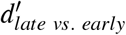 as:

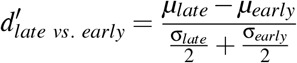

and selected the reward-prediction neurons with 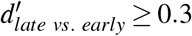. The activity of the reward-prediction neural population in Figure 5 (except Figure 5d) was acquired following *k* -fold cross-validation. We randomly split all trials into 10 folds. we used 9 folds as training trials to compute 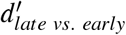. Trial-by-trial activity for the remaining 1 fold (test trials) was computed by averaging across the reward-prediction neurons that met the selection criteria. We repeated this 10 times until average population activity for every fold (and thus every trial) was acquired.

To obtain reward-prediction activity aligned to the first lick (Figure 5j), only rewarded trials (leaf1) with a first lick happening after 0.5 m from corridor entry were included to enable us to investigate the reward prediction signal before licking starts. Due to this criteria, one mouse was excluded because there were no trials with a first lick later than 0.5 m.

### Statistical tests

We performed paired t-tests in Figure panels 1j, 2fj, 3bc, 4di, 5g, S1cd, S3b, S4d, and S5ac; and performed independent t-tests in Figure panels 3eh, 5kl, and 6f. * denotes *p <* 0.05, ** denotes *p <* 0.01, and *** denotes *p <* 0.001. Error bars on all figures represent s.e.m. The exact p-values are below for each figure. Where 4 values are reported, these are for V1, medial, lateral and anterior regions.

- 1j. supervised: 0.940 / 0.00726 / 0.0341 / 0.0261; unsupervised: 0.212 / 3.21*×*10^−4^ / 0.245 / 0.0202
- 2f. supervised: 3.94*×*10^−6^ / 1.72*×*10^−4^ / 1.16*×*10^−5^/ 1.33*×*10^−4^; unsupervised: 1.79*×*10^−8^ / 4.42*×*10^−5^ / 2.22*×*10^−5^ / 2.72*×*10^−5^; naive: 2.83*×*10^−7^ / 6.21*×*10^−4^ / 3.38*×*10^−6^ / 2.19*×*10^−5^
- 2j. supervised: 0.002 / 2.84*×*10^−4^ / 0.00205/ 0.00272; unsupervised: 1.45*×*10^−4^ / 5.22*×*10^−5^ / 1.74*×*10^−4^ / 0.00484; naive: 7.76*×*10^−4^ / 1.40*×*10^−4^ / 2.24*×*10^−5^ / 6.17*×*10^−4^
- 3b. supervised: 0.00375 / 0.0922 / 0.00262 / 0.0136; unsupervised: 2.34 *×*10^−5^ / 0.00809 / 2.96 *×* 10^−4^ / 0.0146
- 3c. 0.0136
- 3e. *P*_*sup. vs naive*_: 0.712/ 7.53 *×* 10^−4^ / 0.221/ 0.0421; *P*_*unsup. vs naive*_: 0.875 / 1.98 *×* 10^−4^ / 0.191 / 0.427; *P*_*sup. vs unsup*_.: 0.861 / 0.236 / 0.924 / 0.0838
- 3h. *P*_*sup. vs naive*_: 0.189 / 0.00573 / 0.0174/ 2.29 *×*10^−4^; *P*_*unsup. vs naive*_: 8.60 *×*10^−4^ / 4.52 *×*10^−5^ / 6.04 *×*10^−3^ / 0.0183 *×*10^−2^; *P*_*sup. vs unsup*_.: 0.147 / 0.259 / 0.593 / 0.00343
- 4d. supervised: 0.00218 / 6.02 *×*10^−4^ / 0.00580/ 0.149; unsupervised: 0.0150 / 0.0106 / 0.00417 / 0.110; naive: 0.518 /0.450 / 0.266 / 0.386
- 4i. supervised:0.00240 / 0.00180 / 0.00346/ 0.202; unsupervised: 5.62 *×*10^−6^ / 1.29 *×*10^−4^ / 9.64 *×*10^−6^ / 0.00437; naive: 7.48 *×*10^−6^ / 0.624 / 0.00169 /0.00619
- 5g. supervised: 0.00690; unsupervised: 0.708
- 5k. 3.41*×*10^−14^
- 5l. 0.995
- 6f. by day: 0.00699 / 0.00447 / 0.0406 / 0.302 / 0.268.
- S1c. (left). supervised: 0.206 / 0.00497 / 0.0297/ 0.0114; unsupervised: 0.0389 / 1.15 *×*10^−4^ /0.982 / 0.249
- S1c. (right). supervised: 0.562 / 0.0219 / 0.104/ 0.134; unsupervised: 0.017 / 0.00130 / 0.0129 / 0.00977
- S1d. (left). supervised: 0.419; unsupervised 0.0466
- S1d. (right). supervised: 0.574; unsupervised 0.126
- S3b. supervised: 0.772 / 0.799 / 0.474/ 0.978; unsupervised: 0.957 / 0.476 / 0.904 / 0.691
- S4d. supervised: 2.06*×*10^−6^ / 5.47*×*10^−4^ / 2.31*×*10^−4^/ 1.41*×*10^−4^; unsupervised: 4.75*×*10^−9^ / 3.30*×*10^−5^ / 1.25*×*10^−7^ / 2.62*×*10^−5^; naive: 4.71*×*10^−10^ / 0.881 / 3.94*×*10^−6^/ 3.33*×*10^−7^
- S5a. supervised: 0.496 / 0.151 / 0.0905 / 0.00690; unsupervised: 0.441 / 0.632 / 0.882 / 0.708
- S5c. supervised: 0.277 / 0.700 / 0.548/ 0.0210; unsupervised: 0.183 / 0.276 / 0.235 / 0.0546

### Retinotopy

Retinotopic maps for each imaging mouse were computed based on receptive field estimation using neural responses to natural images (at least 500 natural images repeated 3 times each). This proceeded in several steps:

1. Obtaining a well-fit convolutional encoding model of neural responses with an optimized set of 200 spatial kernels, using a reference mouse (Figure S1a).
2. Fitting all neurons from our imaging mice to these kernels to identify the preferred kernel and the preferred spatial position (Figure S1b).
3. Aligning spatial position maps to a single map from the reference mouse.
4. Outlining brain regions in the reference mouse using spatial maps and approximately following the retinotopic maps from [30].

Compared to previous approaches, ours takes advantage of single neuron responses rather than averaging over entire local populations, and by using natural images we can better drive neurons and obtain their specific receptive field models. The mapping procedure was sufficiently efficient that it could be performed in a new mouse with responses to only 500 test images each repeated 3 times. Below we describe each step in detail:

**Step 1**. Using the reference mouse, we used the following to model the response of neuron *n* to image img:

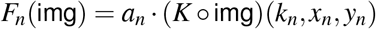

where *a*_*n*_ is a positive scalar amplitude, *º* represents the convolution operation, *x*_*n*_, *y*_*n*_ represent the position in the convolution map for neuron *n* and *k*_*n*_ represents the index of the convolutional map, and finally *K* is a matrix of size 200 by 13 by 13 containing the convolutional filters. This model was fit to neural responses to a natural image dataset of approximately 5,000 images shown at a resolution of 120 x 480, which were downsampled to 30 x 120 for fitting. The kernels *K* were initialized with random gaussian noise. An iterative EM-like algorithm was used to optimize the kernels, which alternated between: 1) finding the best position (*x*_*n*_, *y*_*n*_) for each neuron *n* as well as the best kernel *k*_*n*_ and the best amplitude *a*_*n*_; 2) optimizing *K* given a fixed assignment of (*x*_*n*_, *y*_*n*_, *k*_*n*_, *a*_*n*_) for each neuron *n*. The first part of the iteration was done in a brute force manner: responses of each kernel at each location for each image were obtained and correlated with the responses of each neuron. The highest correlated match for each neuron was then found and its corresponding (*x*_*n*_, *y*_*n*_, *k*_*n*_, *a*_*n*_) were used to fit *K*. The best estimate for kernels *K* was approximately equivalent to averaging the linear receptive field all cells *n* assigned to a kernel *k*_*n*_ after alignment to their individual spatial centers *x*_*n*_, *y*_*n*_. After each iteration, the kernels were translated so their centers of mass would be centered in the 13x13 pixel frame. The center of mass was obtained after taking the absolute value of the kernel coefficients. After less than 10 iterations, the kernels converged to a set of well-defined filters (Figure S1a).

**Step 2**. After the kernels *K* were estimated once, for a single reference recording, we used them for all recordings by repeating the first step of the iterative algorithm in step 1 with a slight modification. Instead of assigning each neuron (*x*_*n*_, *y*_*n*_, *k*_*n*_, *a*_*n*_) independently, we averaged the 2D, maximum correlation maps of the nearest 50 neurons to each neuron, and then took their maximum. This essentially smoothed the spatial correlations to ensure robust estimation even for neurons with relatively little signal (Figure S1b).

**Step 3**. To align spatial maps to the reference mouse, we used kriging interpolation to find a tissue-to-retinotopy transformation *f* . Intuitively, we want to model the data from the alignment mouse as a smooth function *f* from a two-dimensional space of (*z,t*) positions in tissue to another two-dimensional space of retinotopic preferences (x,y). For a new mouse with tissue positions (*z*′,*t*′) and retinotopic positions (*x*′, *y*′), we can then optimize an affine transform *A* composed of a 2x2 matrix *A*_1_ and 1x2 bias term *A*_2_ such that

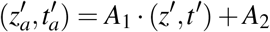

so that

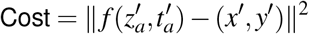

is minimized. To fit the smooth function *f*, we use kriging interpolation, so that *f* is the kriging transform

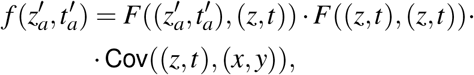

where *F* is a squared exponential kernel *F*(*a, b*) = exp (−∥*a b*∥ ^2^*/*σ^2^) with a spatial constant σ of 200*μ*m and Cov is the covariance between inputs and outputs. Note we can precompute the second part of *f* since it does not depend on 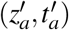. We then optimize the affine transform *A. A* is initialized based on a grid search over possible translation values within *±*500*μ*m. After the grid search, we use gradient descent on the values of *A*, allowing for translation and rotation, but with a regularization term on *A*_1_ to keep the matrix close to the identity. Finally, for some sessions the optimization did not converge, in which case we restricted the matrix *A*_1_ to a fixed determinant, thus preventing a scaling transform.

**Step 4**. The final step was to delineate area borders on the reference mouse, which were then transformed to all mice as described in step 3. Similar to [30] we computed the sign map and parcellated it into regions where the sign did not change. Ambiguities in the sign map were resolved by approximately matching areas to the data from [30]. Note that the exact outlines of the areas in some cases had different shapes from those in [30]. This is to be expected from two sources: 1) the maps in [30] are computed from widefield imaging data, which effectively blurs over large portions of cortex thus obscuring some boundaries and regions; 2) our specific cranial windows are in a different position from [30]. Nonetheless, we do not think the small mismatch in area shapes would have a large impact on our conclusions, given that we combine multiple areas into large regions.

---

**S1:**
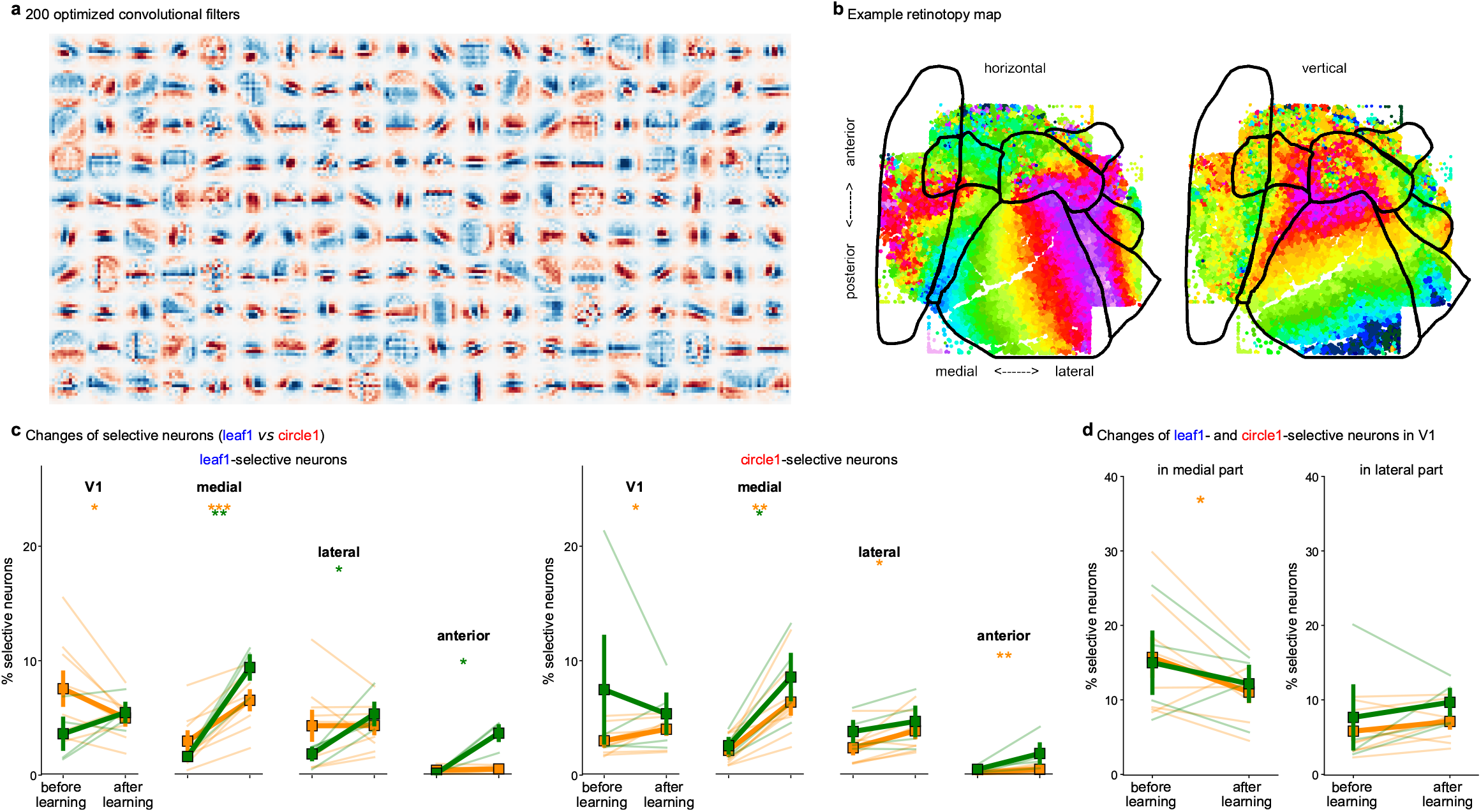
Retinotopy and neural changes after learning for different populations. **a**, Convolutional filter bank used for mapping retinotopy. **b**, Retinotopic maps for an example mouse after alignment to a reference atlas. **c**, Same as Figure 1j split into leaf1-selective (left) and circle1-selective neurons (right). **d**, Similar to Figure 1j for V1 neurons split into a medial and a lateral part.

**S2:**
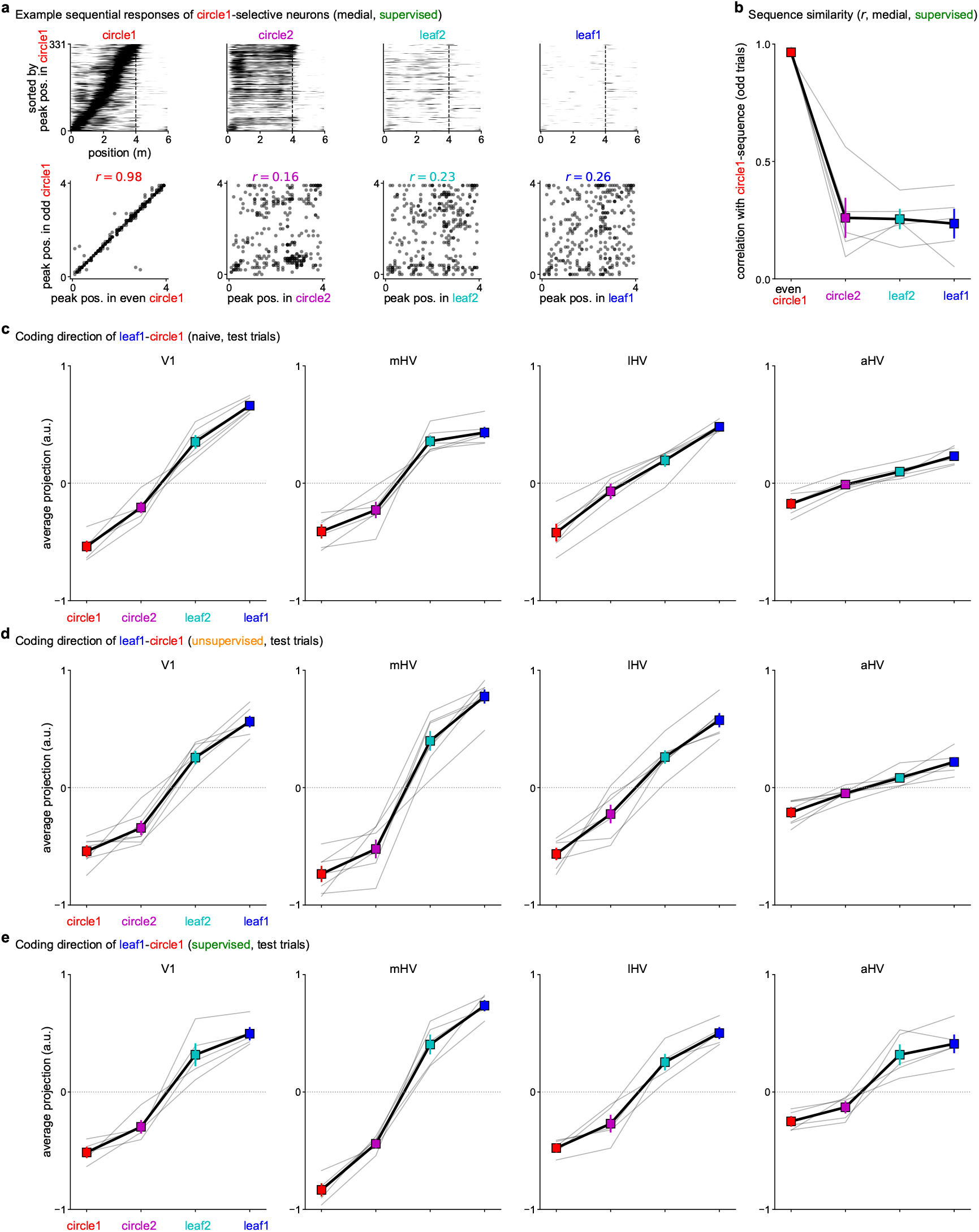
Sequences in circle1-preferring neurons and average projections on the coding direction. **a**, Same as Figure 2d for circle1-selective neurons sorted by the circle1 trials. **b**, Same as Figure 2e for circle1-selective neurons. **c-e** Same as Figure 2i for all brain regions and all mouse cohorts: **c**, naive (n=5 mice, 6 sessions), **d**, unsupervised (n=7 mice), **e**, supervised (n=5 mice).

**S3:**
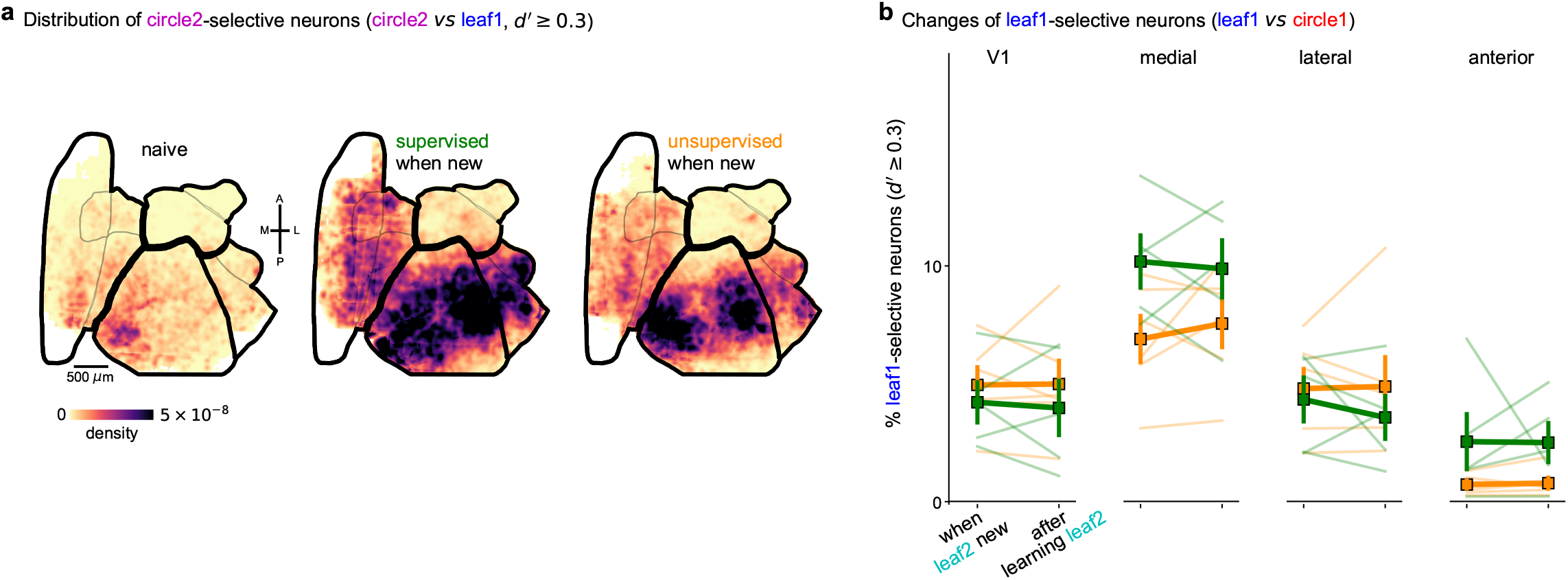
Representations of familiar versus novel stimuli. **a**, Same as Figure 3a for circle2-selective neurons. **b**, Same as Figure 3b for leaf1-selective neurons.

**S4:**
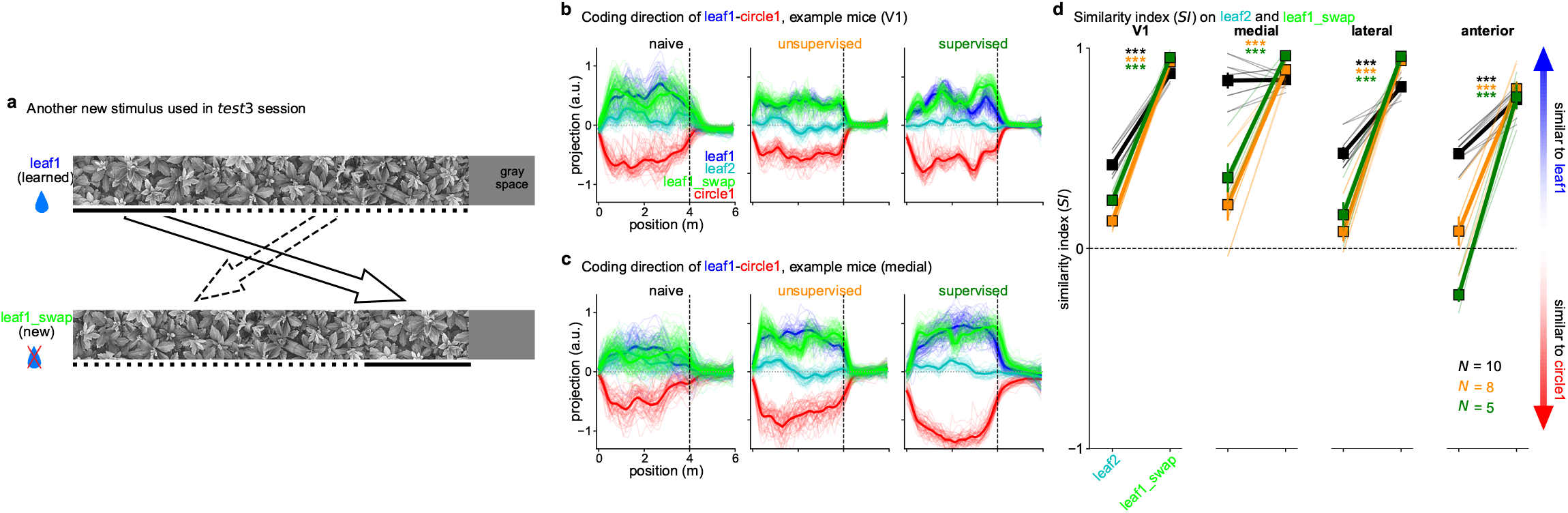
Coding direction projections and similarity indices during *test3*. **a**, A second swap type used in the *test3* session. **b**, Example coding direction of leaf1 vs circle1 in the *test3* session in V1. **c**, Same as **b** in the medial region. **d**, Similarity index along the leaf1-circle1 coding direction in the *test3* session.

**S5:**
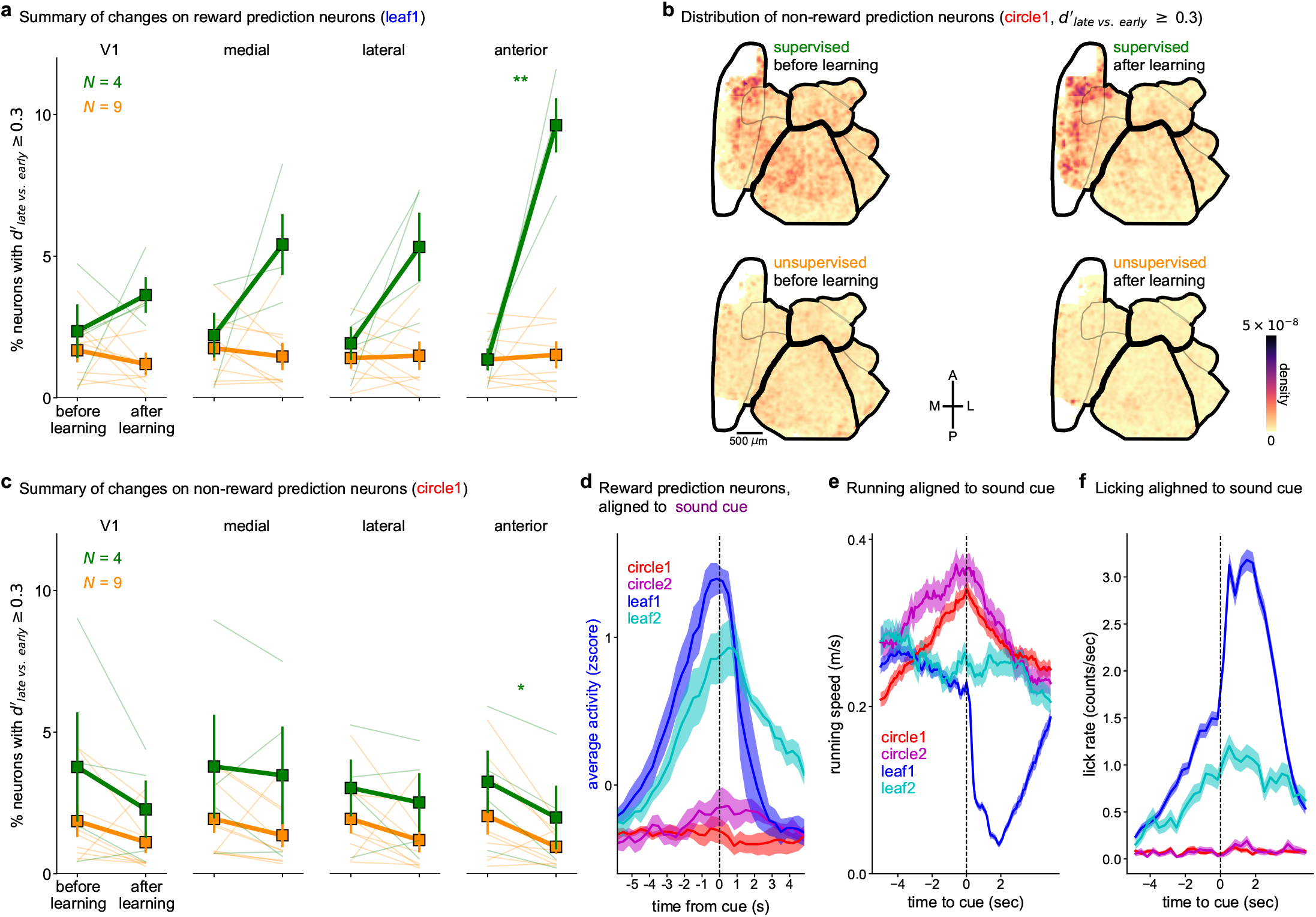
Reward and non-reward prediction neurons across areas. **a**, Percentage of reward-selective neurons before and after learning across all regions. **b**, Distribution of non-reward prediction neurons, defined as neurons with a 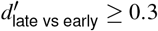 computed from the circle1 corridor trials, which are non-rewarding. **c**, Summary of changes from before to after learning for the non-reward prediction neurons. **d**, Reward prediction neuron average response aligned to the sound cue for all four corridors in the *test1* session. **e**,**f**, Running speed and licking rate respectively, aligned to sound cue.

**S6:**
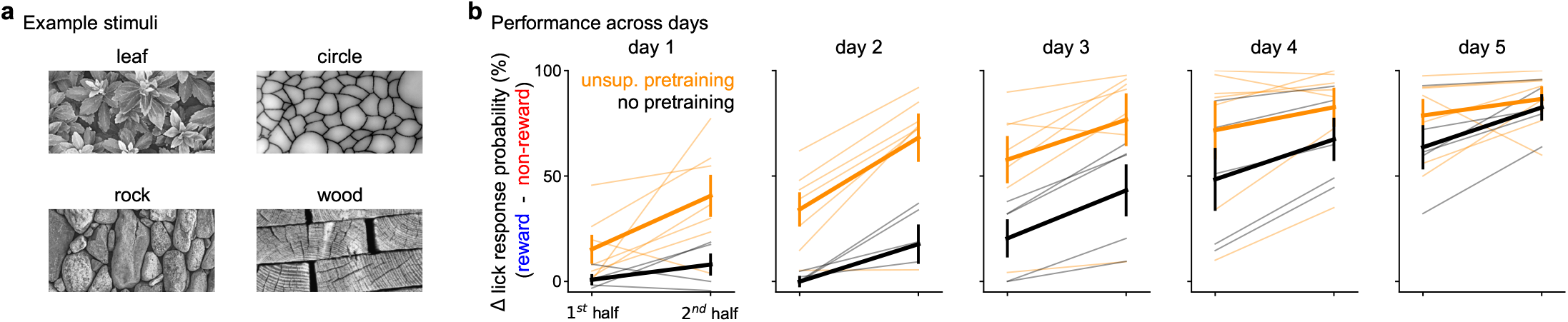
Stimuli for unsupervised pretraining and within-day learning. **a**, Example stimulus crops used for unsupervised pretraining experiments. All pairs of textures were used with the rewarding texture varying from mouse to mouse. **b**, Behavioral performance over days, split into the first and second halves of the session.

